# Microbial diversity characterization of seawater in a pilot study using Oxford Nanopore Technologies long-read sequencing

**DOI:** 10.1101/2020.01.08.898312

**Authors:** M. Liem, A.J.G. Regensburg-Tuïnk, C.V. Henkel, H.P. Spaink

## Abstract

Currently the majority of non-culturable microbes in sea water are yet to be discovered, Nanopore offers a solution to overcome the challenging tasks to identify the genomes and complex composition of oceanic microbiomes. In this study we evaluate the utility of Oxford Nanopore Technologies (ONT) sequencing to characterize microbial diversity in seawater from multiple locations. We compared the microbial species diversity of retrieved environmental samples from two different locations and time points. With only three ONT flow cells we were able to identify thousands of organisms, including bacteriophages, from which a large part at species level. It was possible to assemble genomes from environmental samples with Flye. In several cases this resulted in >1 Mbp contigs and in the particular case of a Thioglobus singularis species it even produced a near complete genome. k-mer analysis reveals that a large part of the data represents species of which close relatives have not yet been deposited to the database. These results show that our approach is suitable for scalable genomic investigations such as monitoring oceanic biodiversity and provides a new platform for education in biodiversity

## INTRODUCTION

Although marine microbes have been studied for multiple decades there is still little knowledge on species diversity in the largest ecological environments of our planet [**1**-**3**]. Current database collections are estimated to represent <5% of oceanic microbial communities [**4**]. Seawater contains many non-culturable organisms, hence to understand its microbial ecology we need to collect sequencing data from DNA samples obtained directly from the environment.

Large-scale metagenomics analyses of seawater have been performed already since 2004 showing remarkable species diversity [**5**]. However, even with availability of abundant sequencing technology resources a complete understanding on the entire diversity remains a challenging task. This is due to, among others, vast water volumes and huge amounts of microbe communities, which through temporal and spatial dynamics contribute to the existence of a near infinite number of ecosystems. Recent studies focussing on marine biodiversity show that a variety of sediments harbour different ecosystems that are particularly extreme in deep ocean environments. There have been many exploratory studies of harnessing marine microorganism for the production of bioactive compounds, with versatile medicinal, industrial, or agricultural applications [**6**].

Microbial diversity characterization has primarily relied on traditional high-throughput short-read sequencing methods, such as Illumina [**7**-**12**] or 454 sequencing [**5**]. Even though Pacific Biosciences single-molecule long-read sequencing has been used to catalogue the diversity of coral-associated microbial communities, these studies relied on amplification and 16S rRNA homology to position microbes taxonomically [**5**,**7**,**13**,**9**-**11**,**14**]. Amplification, however, introduces biases that results in over- and underrepresentation of particular species. Additionally, in some cases 16S rRNA identification fails to characterize microbial diversity due to variability in the 16S region [**15**], - for example, previous studies revealed that some universal primers have strong biases against the detection of pelagic bacteria (SAR11 group) and archaea [**4**]. Hence 16S-based methods appear ineffective at comprehensively characterizing complex metagenomics samples such as from seawater. Furthermore, traditional 16S rRNA identification is limited to the detection of microbe presence and does not yield further functional insights about the organism. And finally, high-throughput short read sequencing methods require large scale infrastructure including sequencers and laboratories.

In this pilot study we evaluate the utility of Oxford Nanopore Technologies (ONT) sequencing to characterize microbial diversity in seawater. ONT sequencing generates on average 10 Kbp reads, theoretically without upper limit, and bypasses the necessity of amplification. Our strategy aims to classify microbial diversification directly from environmental samples (two different oceanic locations were chosen) with minimal computational and financial cost over a relatively short time span. This will facilitate future scalable investigations such as monitoring oceanic biodiversity and the time and space dynamics these microbes are subject to.

## RESULTS

### Sample collection, data quality control and verification of microbial content

We collected samples from coastal regions of both the Atlantic Ocean (west part of the English Channel - Roscoff, France, August 2017) and the south part of the North Sea (Wassenaarseslag, the Netherlands, July 2017 and August 2018). From here on, we refer to these as samples 1, 2 and 3. MinION 48-hour sequencing runs on every sample resulted in three datasets with mean read lengths that range between 1,511 and 7,983 bp (Table 2). Our read length distributions indicate relatively suboptimal DNA samples that resulted in shorter reads (Figure 1) compared to ONT read length averages of laboratory cultures. This is particularly apparent for sample 1. The error rate expressed in PHRED indicates similar quality for the three runs, our average qualities fluctuate around PHRED 12 that stands for <10% error per read on average.

**Table 2.** Raw sequencing data statistics of sample 1,2 and 3

| Statistics | France (1) | The Netherlands '17 (2) | The Netherlands '18 (3) |
| --- | --- | --- | --- |
| Reads | 370,371 | 1,316,823 | 225,200 |
| Bases | 559,696,414 | 6,350,530,291 | 1,797,851,809 |
| Mean length (bp) | 1,511 | 4,822 | 7,983 |
| Max length (bp) | 49,807 | 155,979 | 161,655 |
| 16S reads | 23 | 178 | 188 |

**Figure 1.**
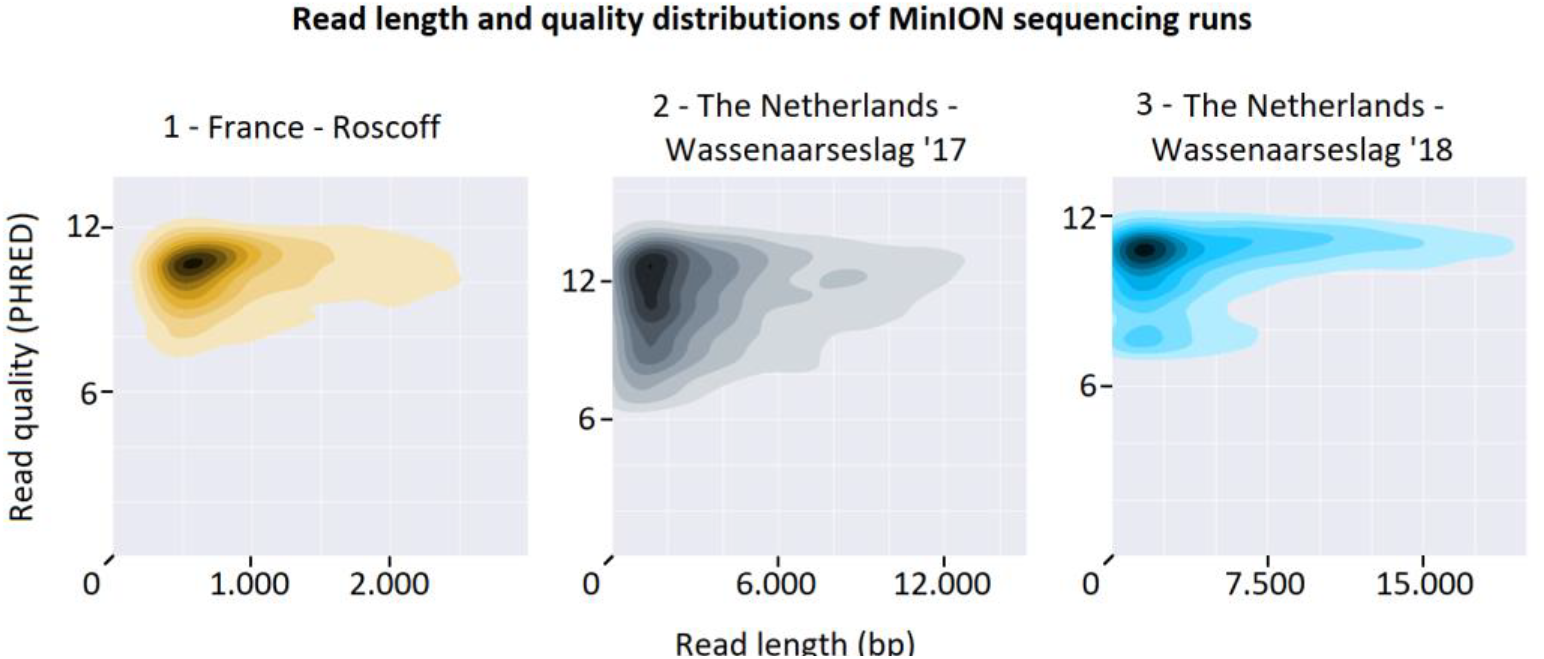
Read length and quality distributions of 48-hour run sequencing data for sample 1, 2 and 3 (from left to right). Mean read lengths vary from 1,511 up to 7,983 bp with similar base call qualities (around PHRED 12). Plots are based on NanoPlot plotting [23]

In order to asses quality of the data we analysed homologues sequences of the three longest reads for all three data sets. The results (Table 1) show that several of these reads are representative of bacterial species that were found to be dominant by the OneCodex analyses. One of the reads (France - Read ID 1) also showed that we have identified a representative of a bacteriophage of Pelagibacter. The limited coverage of the homologues gene indicates that we have identified a rather distant new relative of the published bacteriophage.

**Table 1.** Blast alignment of longest raw sequencing reads. **Sample)** time and ocation of seawater samples, **Read ID)** read length identifier sorted from longest to smallest, **Query length)** the length of the read, **Best hits*)\*** criteria for best hit; largest query coverage with highest identity and published study, **Cov)** alignment percentage that reads cover the reference, **ID)** alignment identity between query and reference, **Ref length)** length of the reference sequence

| Sample | Read ID | Query length (Kbp) | Best hits * | Cov (%) | ID (%) | Ref length (Kbp) |
| --- | --- | --- | --- | --- | --- | --- |
| Fr. | 1 | 50 | Pelagibacter phage HTVC008M [29] | 2 | 78 | 147 |
| Fr. | 2 | 46 | Candidatus Pelagibacter sp. FZCC0015 [CP031125] | 3 | 68 | 1,364 |
| Fr. | 3 | 45 | Halioglobus pacificus strain RR3-57 [CP019046] | 9 | 69 | 4,847 |
| NL '17 | 1 | 155 | Brassica oleracea HDEM [LR031920] | 3 | 68 | 113 |
| NL '17 | 2 | 107 | No hit –repetitive stretch |  |  |  |
| NL '17 | 3 | 78 | Halioglobus japonicus strain NBRC 107739 [CP019450] | 2 | 69 | 4,085 |
| NL '18 | 1 | 161 | Flavobacterium columnare [31] | 28 | 68 | 3,329 |
| NL '18 | 2 | 149 | Clostridium tetani strain Harvard 49205 [CP035787] | <1 | 69 | 2,807 |
| NL '18 | 3 | 139 | Micromonas sp. RCC1109 virus MpV1 [30] | 23 | 74 | 184 |

To confirm that our double filtering method indeed selects for microbial DNA we have used 16S rRNA primers that are known to identify a wide range of microbial genomes. FastPCR aligns the currently ‘best available’ 16S rRNA primer sequences **[25]** to raw sequencing data and shows microbial content in all three raw sequencing datasets. We found 23, 178 and 188 hits aligning both forward and reverse primers that span between 420 and 470 bp (Table 2). These hits have a minimum of 80% alignment identity and ranged up to 100% matches. Blast searches of regions that have <80% sequence identity did not result in hits originating from 16S rRNA hence do not contribute to the identification of microbial content and have been omitted.

### Seawater characterization using k-mer classification

Using OneCodex **[26]** we generated classification trees for the three datasets. These are built from raw sequencing data and indicate the taxonomic relation between the detected microbial classes. This relation is based on taxonomic identifiers (taxids) provided by the NCBI taxonomy database. For visualization purposes these taxonomic trees are subsets of the complete classifications: every node is supported with a minimum threshold of 831, 2,048 and 588 reads for samples 1, 2 and 3, respectively.

Despite the fact that a large part of all three datasets could not be classified (47%, 69% and 38% for sample 1, 2 and 3, respectively), all taxonomic trees highlight the complexity of microbial communities present at a single site. None of our three datasets reveal an overall dominant species, with the largest differences between samples microbes that appear at low abundances. However 4.46% (sample 1), 15.66% (sample 2) and 7.82% (sample 3) of classified reads belong to Planktomarina temperata, which is therefore the most abundant species present in the three data sets combined (Figure 2, Figure 3 and Figure 4, red nodes).

**Figure 2.**
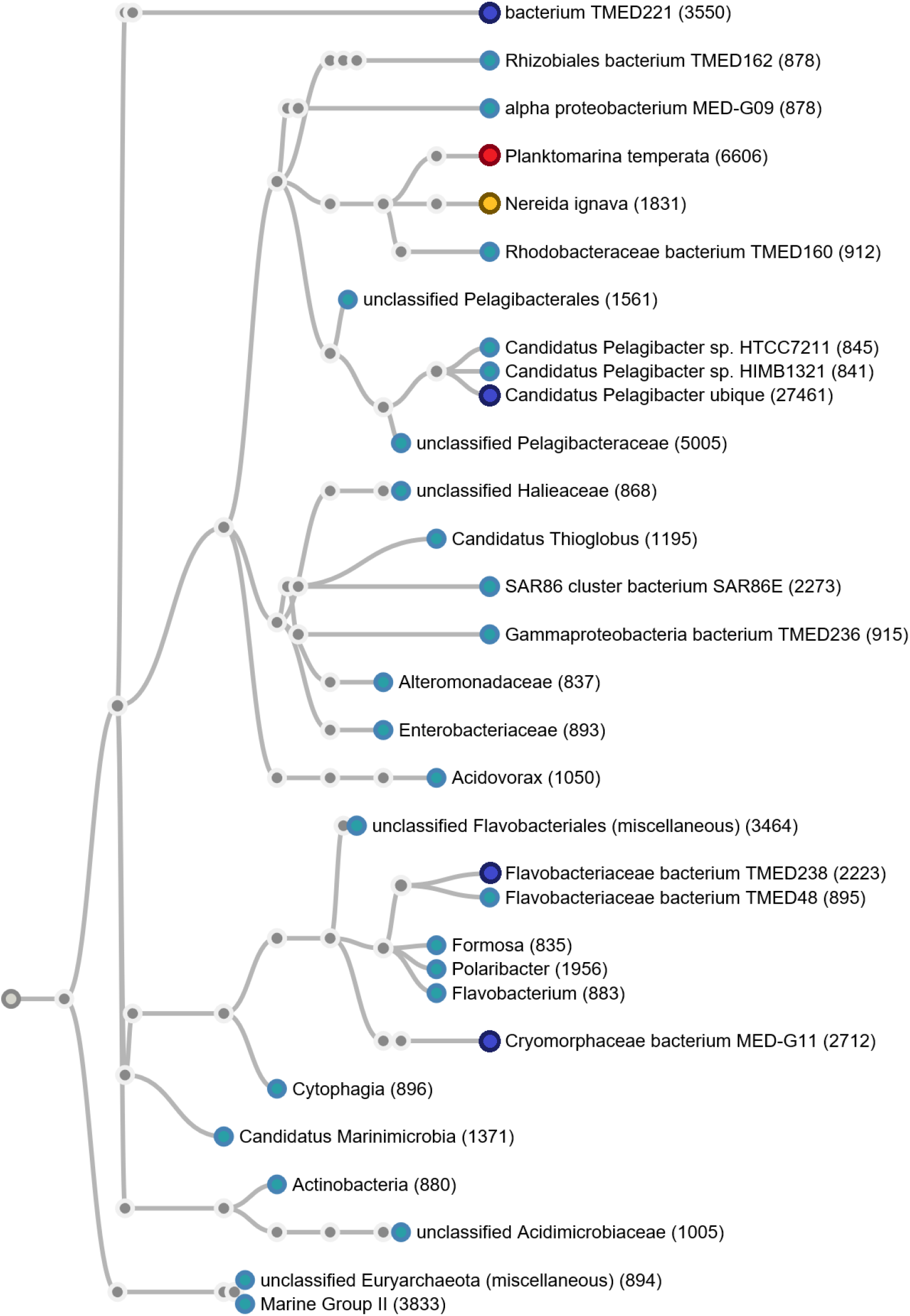
Taxonomic tree on a subset of the data generated from sample 1 data. Every node stands for a taxonomical ID that is supported with at least 831 reads. In red the most abundant species present in all three samples. Dark blue nodes together with the red node highlight the top-5 most abundantly present species in this sample. The yellow node indicates the most prominent species difference between the two locations.

**Figure 3.**
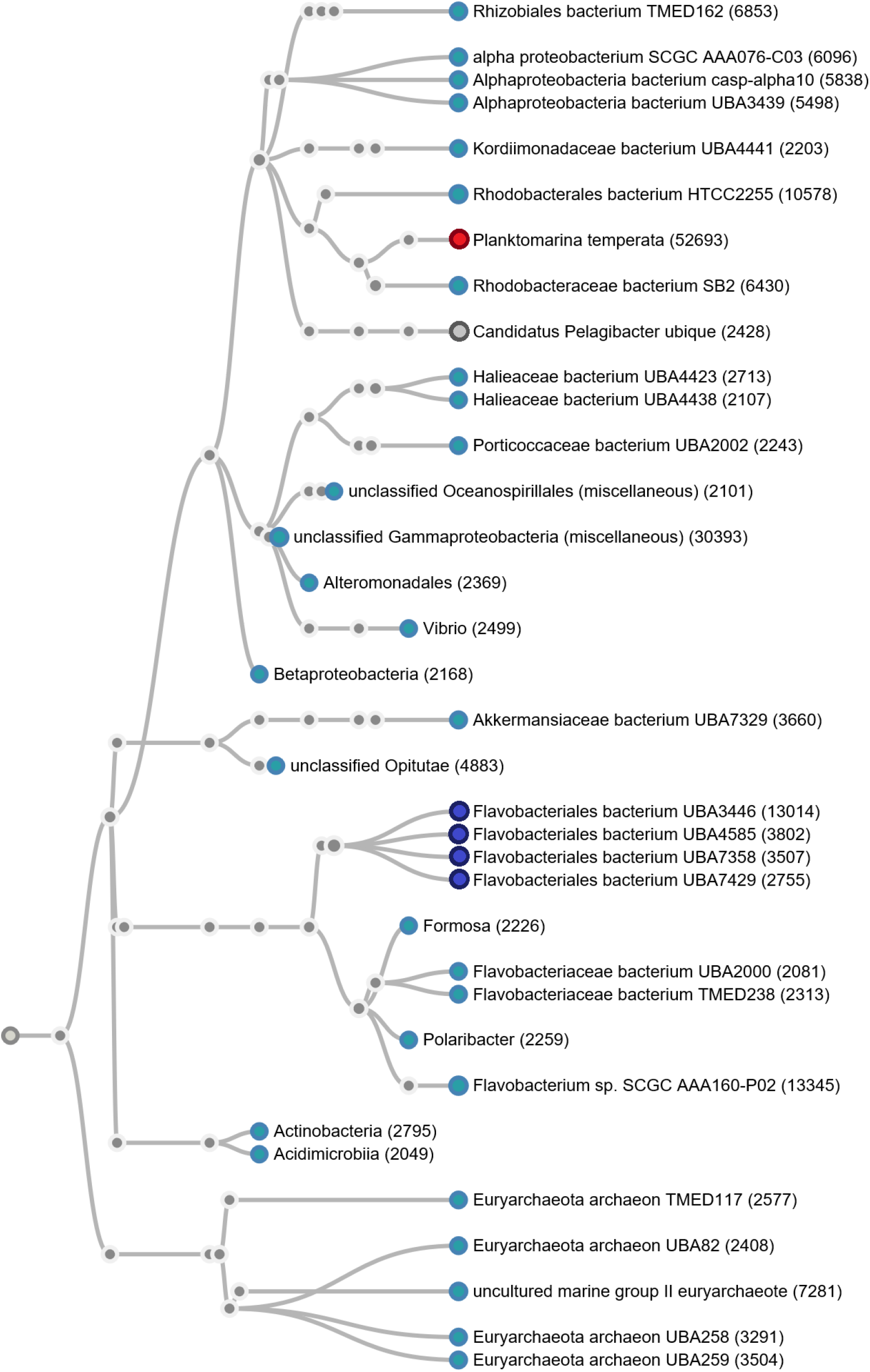
A subset of the data set from sample 2, every node is supported with minimally 2048 reads. The red node indicates the most abundant species over all three datasets, together with dark blue nodes it comprises the top-5 most abundant species in this dataset. Particularly underrepresented is species Candidatus Pelagibacter (grey node) compared to sample 1 and 3.

**Figure 4.**
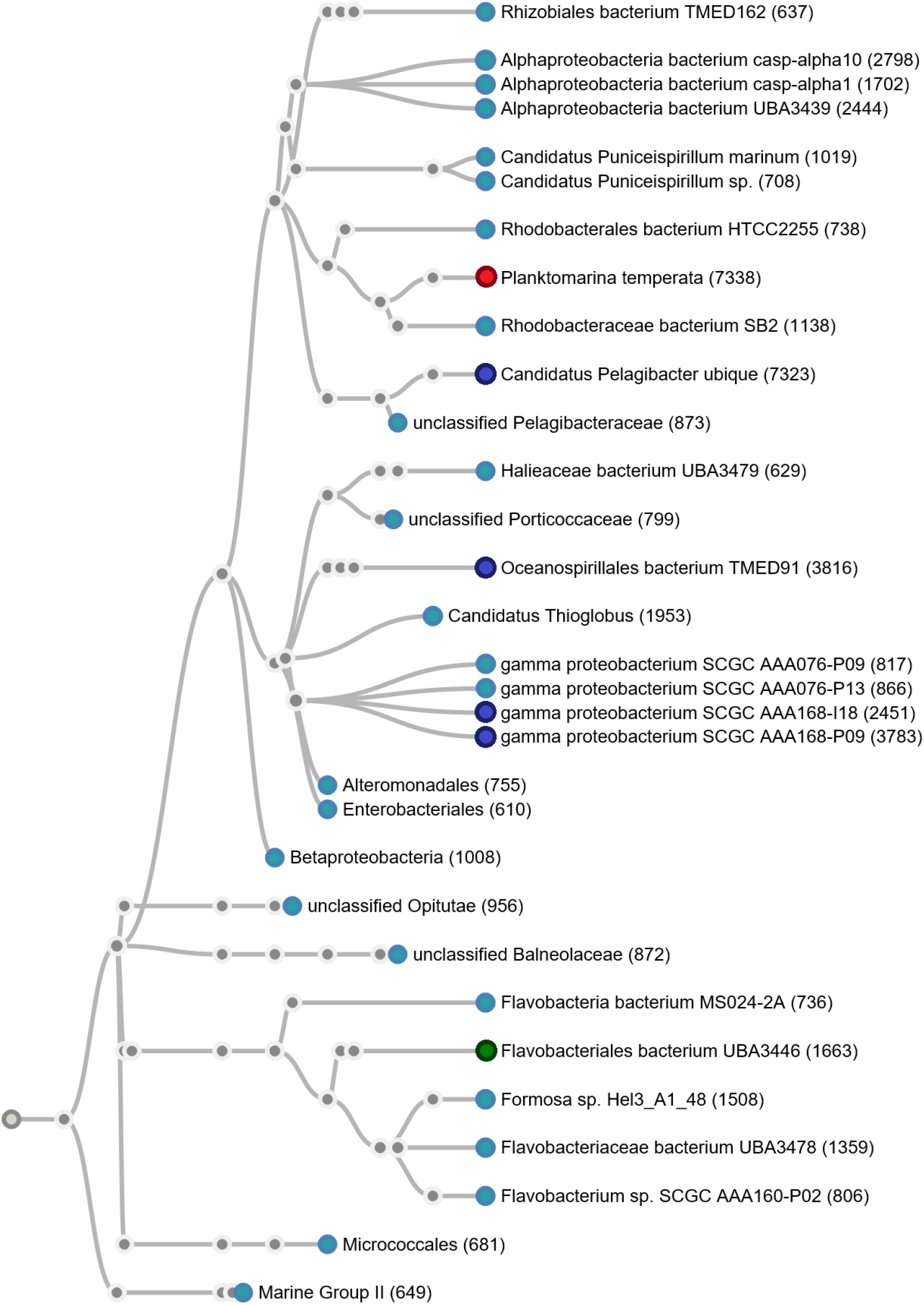
Taxonomic tree on a subset of sequencing data from sample 3, every node is supported with at least 588 reads. Again the red node indicates the overall most abundant species, and together with dark blues nodes they form the top-5 most abundant species for this dataset. Compared to the year before Flavobacteriales bacterium is underrepresented (green node).

The top-5 most abundant species in sample 1 are: Candidatus Pelagibacter ubique (9.31% of Proteobacteria), bacterium TMED221 (8.61% of unclassified bacteria), Flavobacteriaceae bacterium TMED238 (4.48% of the FCB group), Planktomarina temperata (4.46% of Proteobacteria) and Cryomorphaceae bacterium MED-G11 (4.16% of the FCB group) (Figure 2, red and dark blue nodes). Approximately 2% of classified reads belong to species Nereida ignava, compared to less than 0.04% from sample 2 and 3 it is the most prominent difference between the two locations (Figure 2, yellow node).

In the second sample four of the top-5 most abundant species belong to the same species: Planktomarina temperata (15.66% of Proteobacteria), Flavobacteriales bacterium UBA3446 (5.54% of the FCB group), Flavobacteriales bacterium UBA7358 (5.30% of the FCB group), Flavobacteriales bacterium UBA4585 (5.12% of the FCB group) and Flavobacteriales bacterium UBA7429 (4.41% of the FCB group) (Figure 3, red and dark blue nodes). Even though Planktomarina temperata reads are abundantly present in all three samples they are particularly enriched (15.66%) in this sample compared to 7.82% from the next year and 4.46% from France. Additionally, the presence of Candidatus Pelagibacter ubique is underrepresented in this sample, 1% of all classified reads belong to this species, compared to ~11% and 9% in sample 1 and 3, respectively (Figure 3, grey node).

Finally, the top-5 most abundant species from sample 3: Candidatus Pelagibacter ubique (9.24% of Proteobacteria), Oceanospirillales bacterium TMED91 (8.12% of Proteobacteria), gamma proteobacterium SCGC AAA168-P09 (8.08% of Proteobacteria), Planktomarina temperata (7,82% of Proteobacteria) and gamma proteobacterium SCGC AAA168-I18 (7.25% of Proteobacteria) (Figure 4, red and dark blue nodes). Interestingly, the species gamma proteobacterium are classified strain specific (Figure 4, dark blue nodes) as opposed to Flavobacteriales bacterium species from sample 2 and is less abundant in this sample (1.6%) compared to the year before (5.9%) (Figure 4, green node).

The taxonomic levels assigned by OneCodex range from kingdom down to species-specific. Reads that cannot be linked to a particular taxonomic level are labelled ‘no rank’. In total 1,750, 3,017 and 2,007 taxids are assigned to the data of sample 1, 2 and 3, respectively. More than half of the ranks that OneCodex was able to classify are assigned to species level (Figure 5 B) in all three samples.

**Figure 5.**
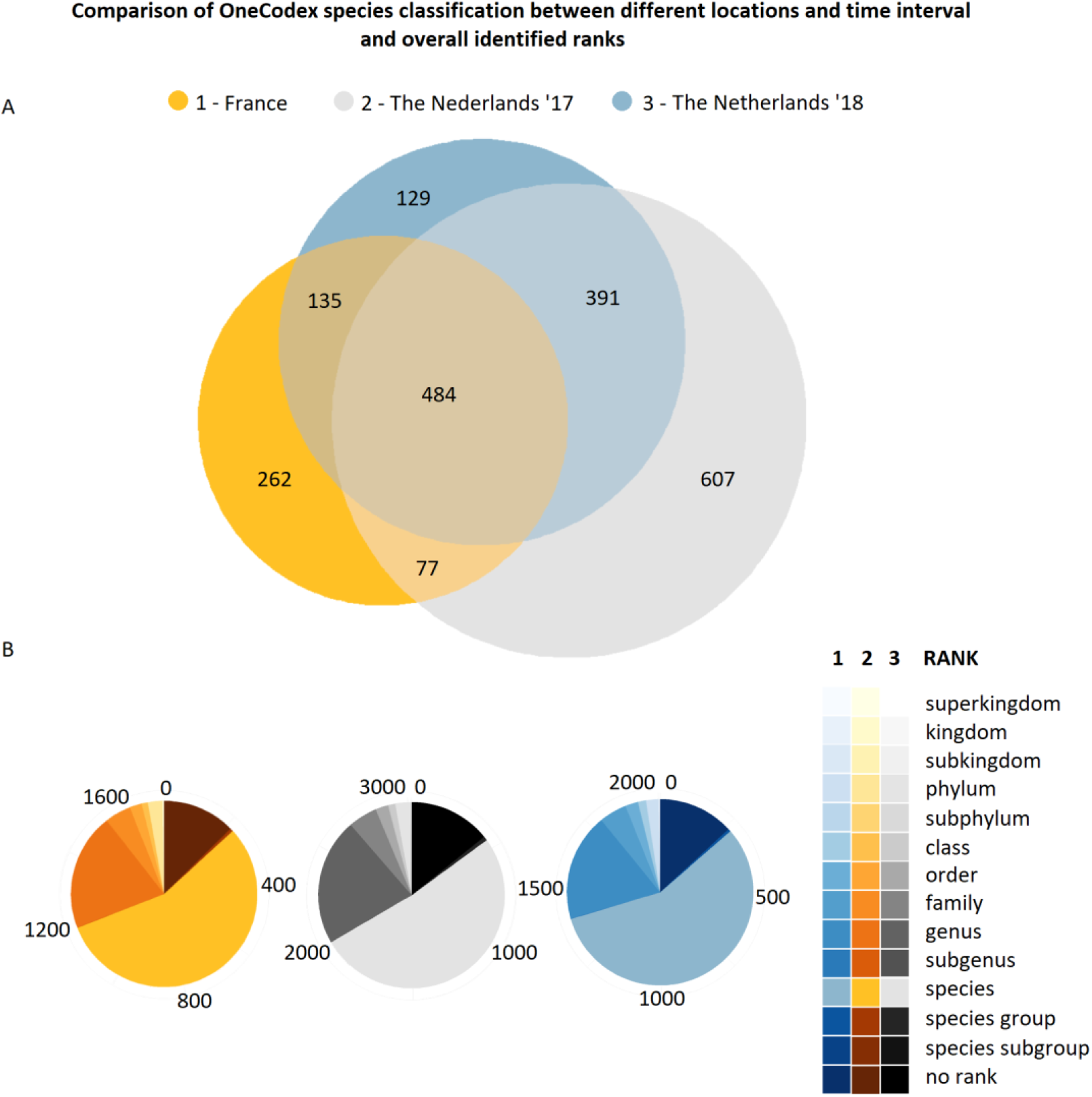
A) Venn diagram ccomparison of identified species by OneCodex, highlighting species that are time and space dependent and also microbes that are not. **B)** Overall OneCodex classification ranks per dataset, the majority of classified reads have been linked to a species

Interestingly, at least 484 microbes are identified in all three samples. Some highlights include: 92 different Flavobacteriaceae bacterium and Flavobacteriales bacterium strains; 19 different Candidatus Pelagibacter strains; 18 Pelagibacteraceae bacterium and 6 SAR strains. This indicates that these communities are less time and location dependent compared to the 262 and 1,127 species that were found exclusively in France or Dutch areas, respectively. Furthermore, 607 and 129 species are exclusively observed in the Netherlands. As they exist at different times, they provide an initial impression of the time-dependent dynamics of these local communities. Finally, 135 and 77 species could be identified that are present at both locations, however only detectable at particular times. This could be an indication that even over large areas microbes are subject to time regulated dynamics.

### Metagenomics assembly on raw sequencing data and blast verification on the top-3 longest contigs

In an attempt to very OneCodex classification results as well as to assess the current metagenomics assemblers capabilities we subsequently assembled the three datasets separately. We have assembled our complex metagenomics datasets with Flye and retrieved 256, 1,735 and 968 contigs with mean coverage of 14x, 13x and 10x from samples 1, 2 and 3, respectively (Table 3). Coverage on contigs ranged up to 62, 89 and 107 for samples 1, 2 and 3, respectively, with a lower-bound of 3x coverage for all three assemblies. As expected, assembly statistics on sample 1 show the least optimal assembly results (lowest number of contigs, smallest mean and max contig lengths and smallest N50 values) given that the data of this sample was smallest in volume with shortest average read lengths. Notably, although it has higher coverage, assembly results from sample 2 did not exceed results from sample 3. On the contrary, sample 3 resulted better average contig length, maximum contig length and N50 values compared to sample 2 (Table 3 and Table 4).

**Table 3.** Flye assembly statistics

| Assembly stats | France (1) | The Netherlands '17 (2) | The Netherlands '18 (3) |
| --- | --- | --- | --- |
| contigs | 256 | 1,735 | 968 |
| length (bp) | 8,678,102 | 107,863,873 | 94,117,952 |
| min length (bp) | 2,432 | 536 | 494 |
| mean length (bp) | 33,898 | 62,169 | 97,229 |
| max length (bp) | 219,363 | 1,098,797 | 1,648,106 |
| N50 | 40,621 | 75,928 | 153,524 |

**Table 4.** Blast alignment for top-3 longest contigs for sample 1, 2 and 3. **ID)** identity number provided by Flye, **Query len)** the length of the contigs, **Cont cov)** data coverage for every contig, **Best hits *)** * criteria for best hit; largest query coverage with highest identity and published study, **Query cov)** how much of the contig covers the reference sequence, **Aln ID)** alignment identity between the reference and contig, **Ref len)** the length of the reference sequence the contig is aligned to.

| Sample | ID | Query len (Kbp) | Cont cov | Best hits * | Query cov (%) | Aln ID (%) | Ref len (Kbp) |
| --- | --- | --- | --- | --- | --- | --- | --- |
| 1 | 23 | 219 | 30 | *Candidatus Pelagibacter ubique* HTCC1062 [17] | 88 | 78 | 1,308 |
| 1 | 227 | 141 | 16 | *Candidatus Actinomarina minuta* [16] | 24 | 82 | 41 |
| 1 | 130 | 137 | 13 | *Candidatus Actinomarina minuta* [16] | 16 | 79 | 36 |
| 2 | 190 | 1,098 | 24 | *Sphingobacterium* sp. EB080_L08E11 [18] | 7 | 93 | 140 |
| 2 | 71 | 1,017 | 26 | marine bacterium Betaproteobacterium [19] | 10 | 94 | 44 |
| 2 | 8 | 967 | 27 | marine bacterium Gammaproteobacterium [20] | 4 | 80 | 61 |
| 3 | 58 | 1,648 | 20 | *Candidatus Thioglobus singularis* [28] | 75 | 80 | 1,714 |
| 3 | 376 | 1,283 | 7 | Uncultured Flavobacteriia bacterium [21] | 4 | 98 | 36 |
| 3 | 206 | 1,138 | 12 | marine bacterium [AY458647] | 4 | 93 | 44 |

Impressively, Flye was able to reconstruct a full genome from our third sample: 75% of our 1.6 Mbp contig aligns with 80% identity to Candidatus Thioglobus singularis of which its complete genome is a single circular chromosome of 1.7 Mbp, with only 20x coverage on this particular contig (Table 4). The longest contig (219 Kbp) assembled from sample 1 represents a fragment of an entire genome and aligns with 88% identity to Candidatus Pelagibacter ubique, from which reads are most abundantly present in sample 1 (Table 4).

Even though OneCodex indicates that only 397 reads originate from Candidatus Actinomarina, Flye was able to reconstruct contigs that exceed the length of the currently available reference sequence. The second (141 Kbp) and third (137 Kbp) longest contigs aligned with 82% and 79% identity to the reference that is just 41 Kbp in size (Table 4). Similarly, Flye results in a top-3 longest contigs from sample 2 and 3 that align with high homology to the reference and all contigs exceed the length of the reference sequence (Table 4).

### Comparison of Flye assembly and raw sequencing data using OneCodex characterization

In order to verify if new species could be identified after assembly we have compared the OneCodex classifications using assembly results to the classification results based on raw sequencing data. Using the 256 contigs Flye was able to reconstruct OneCodex identified 41 species in total from sample 1 (Figure 6). Since reads that originate from Flavobacteriaceae and Pelagibacteraceae are represented in high abundance it is no surprise that detailed species-level classification for these two families appeared most effective, into 9 and 12 strains (out of the 41 classified species), respectively. OneCodex is able to identify 12 species only after assembly, these include 11 deferent Pelagibacteraceae bacterium strains and a SAR86 strain.

**Figure 6:**
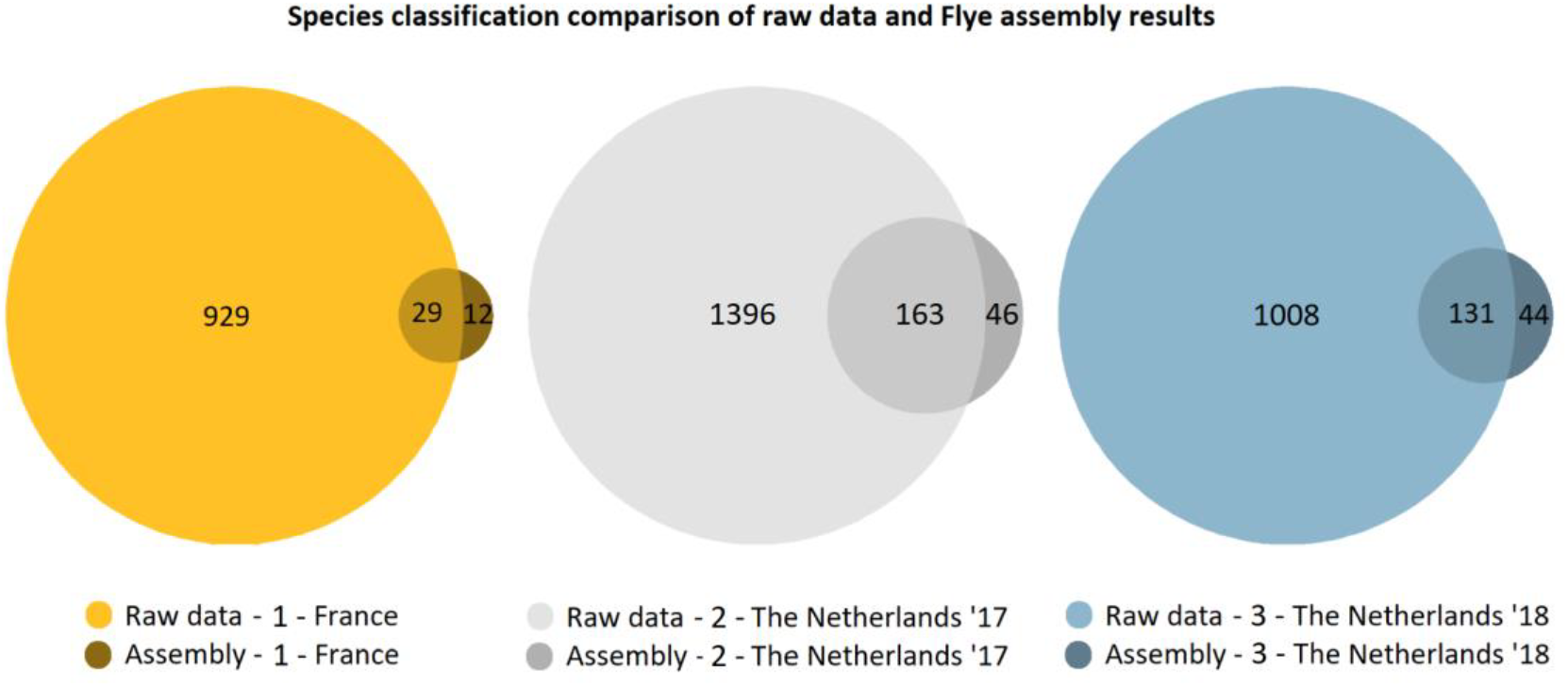
Species classification on sample 1,2 and 3. Lighter shades indicate identified species on raw sequencing data, darker shades highlight species only identifiable after assembly.

Although OneCodex was able to identify the most species using assembly results of sample 2, no prominent strain-specific enrichment was observed exclusively for assembly results this sample. From the 209 species that are identified Flye favoured 5 species during assembly: Alphaproteobacteria bacterium (10 strains), Euryarchaeota archaeon (15 strains), Flavobacteriaceae bacterium (23 strains), Flavobacteriales bacterium (18 strains) and Gammaproteobacteria bacterium (19 strains).

Species diversification of assembly results from sample 3 appeared best for 14 different Flavobacteriaceae bacterium strains, 13 gamma proteobacterium strains, and 13 strains of Gammaproteobacteria bacterium. Notably, 6 Pelagibacteraceae bacterium strains could be identified using assembly results, that could not be classified based on raw sequencing data alone.

### Data quality of unclassified reads and additional in silico PCR analysis

Poor read quality and relatively short read lengths could be a potential reason explaining why OneCodex was unable to classify taxids. Therefore, we investigated quality and length of unclassified reads (Figure 7). Although average lengths are shorter, and average quality values have a larger distribution, the differences are minimal compared to raw sequencing data (Figure 7). These statistics indicate that, in theory, the reads should provide OneCodex with sufficient information to resolve classifications. That OneCodex was not able to classify these reads even to the most general taxonomic levels (such as kingdom or phylum) adds to the notion that these reads originate from species that are novel.

**Figure 7:**
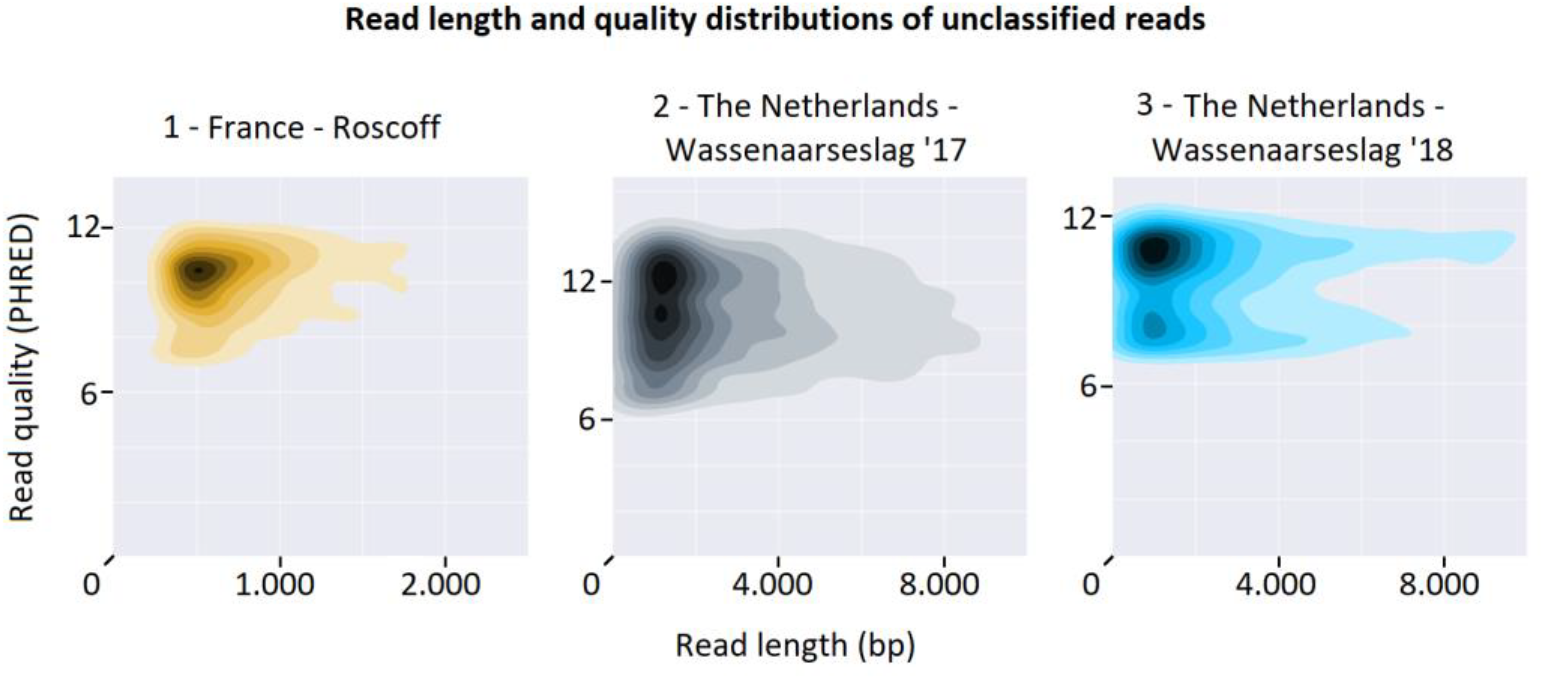
Read length and quality distributions of data that OneCodex labels unclassified. On average reads are shorter compared to raw sequencing data, however these lengths should still be sufficient to use for k-mer species characterization. Average quality distributions are very comparable to reads which OneCodex was able to classify species with.

The proportion of reads for which no classification could be assigned ranges between 38% and 69% compared to the raw sequencing data (Table 5) and provides a general impression on the amount of potentially novel microbes that thrive in these waters. Since OneCodex is particularly tailored to the identification of microbial DNA, unclassified reads potentially belong to non-microbial organisms. We therefore performed an additional round of in silico PCR analysis to inspect the presence of any remaining microbial 16S rRNA fragments. Interestingly, we found at least 10 more reads in sample 2 that have over 80% homology with our primers, showing that microbial content still exists within these unclassified reads (Table 5).

**Table 5.** Data statistics on reads for which OneCodex could not resolve any classification

| Stats | France<br>Unclassified (1) | The Netherlands '17<br>Unclassified (2) | The Netherlands '18<br>Unclassified (3) |
| --- | --- | --- | --- |
| reads | 172,843 | 908,744 | 86,653 |
| bases | 214,536,717 | 3,517,616,897 | 479,777,298 |
| mean length | 1,241 | 3,870 | 5,536 |
| max length | 33,295 | 155,979 | 132,486 |
| % of original data | 47 | 69 | 38 |
| 16S occurrence | 0 | 10 | 0 |

### Inspection of low complexity regions in unclassified reads using tandem repeat analysis

An additional circumstance that might explain why reads are left unclassified is the presence of low complexity regions such as repeat elements. These elements cause k-mers to contain the exact same genomic content making it impossible to assign them uniquely to specific taxa or even to a more general taxonomic level. We have analysed the presence of repeat elements with Tandem Repeat Finder [22] in raw sequencing data and compared these to repeat counts of the unclassified reads. In none of our samples did we observe an increased presence of repetitive elements, on the contrary, the repetitive element count is lowered in every case (Figure 8).

**Figure 8:**
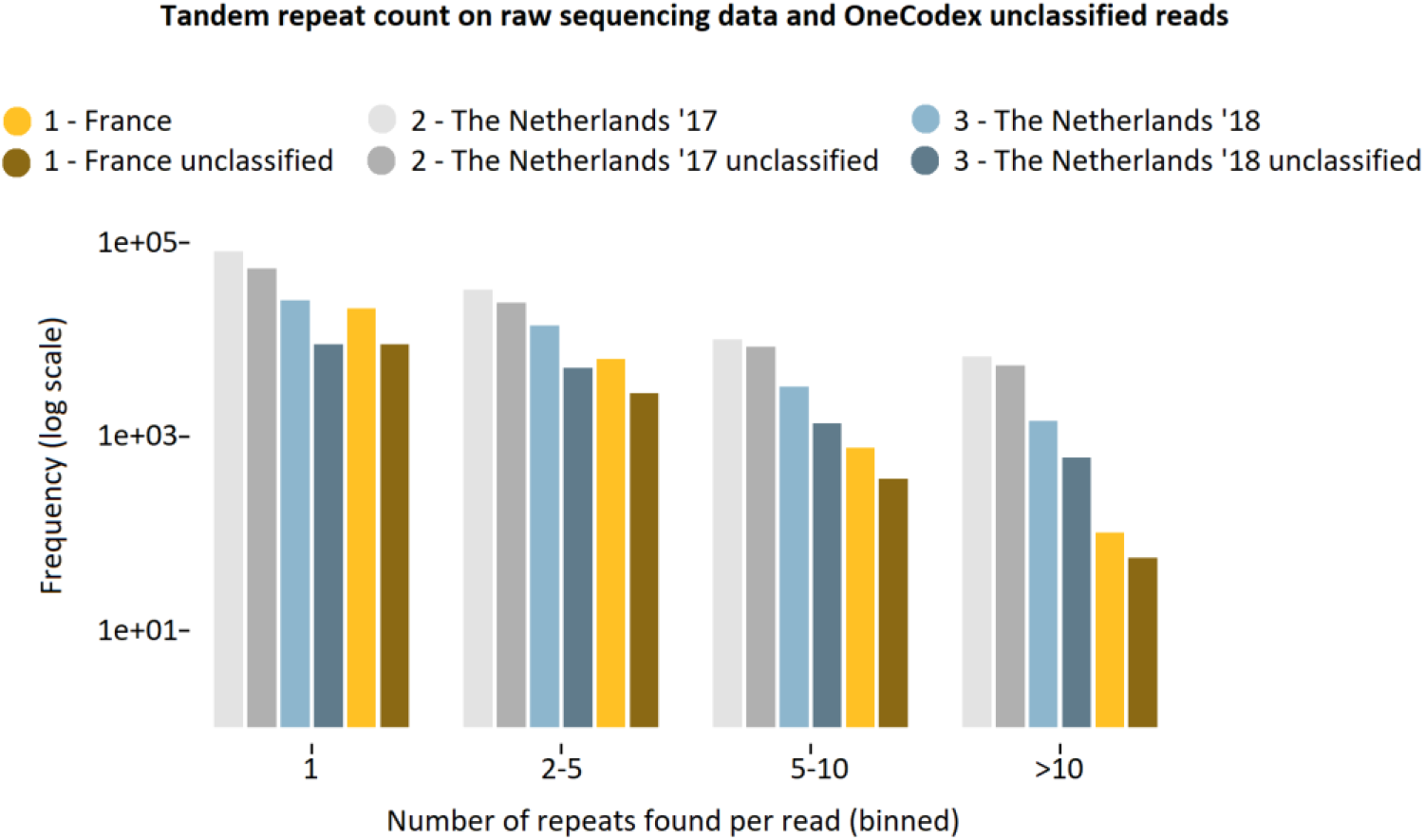
Tandem repeat analysis, counts per read and comparison between raw sequencing data and unclassified data set for different locations and time. Repeat counts are represented in bins, the bins indicate the number of occurrences per read.

Taking together the data characteristics and the lack of both general taxonomical classification and highly abundant regions of low complexity suggest that these reads indeed originate from novel species. It highlights at least the absence of these species in currently publicly available OneCodex database, and provides a general glimpse of the amount of unknown species that comprise oceanic microbiomes.

## MATERIALS AND METHOD

### Sample collection and DNA isolation from salt water

Approximately 10 liter salt water of both locations was filtered through a double filter setup (Figure 9 A). 1.2 μm and 0.22 μm filters are used to remove eukaryotes and phages/ viruses from the samples, respectively (Figure 9 B). Water is passed through a 1.2 μm filter that aims to capture eukaryotic cells on top and is discarded, the remaining water is passed through a 0.22 μm filter during a second filtering round. The microbial material is captured on top of the filter, water that passed through this filter contains phage/viral material and is discarded afterwards. Material captured on the 0.22 μm filter represents the microbiome of our sample and was used for cell lysis.

**Figure 9.**
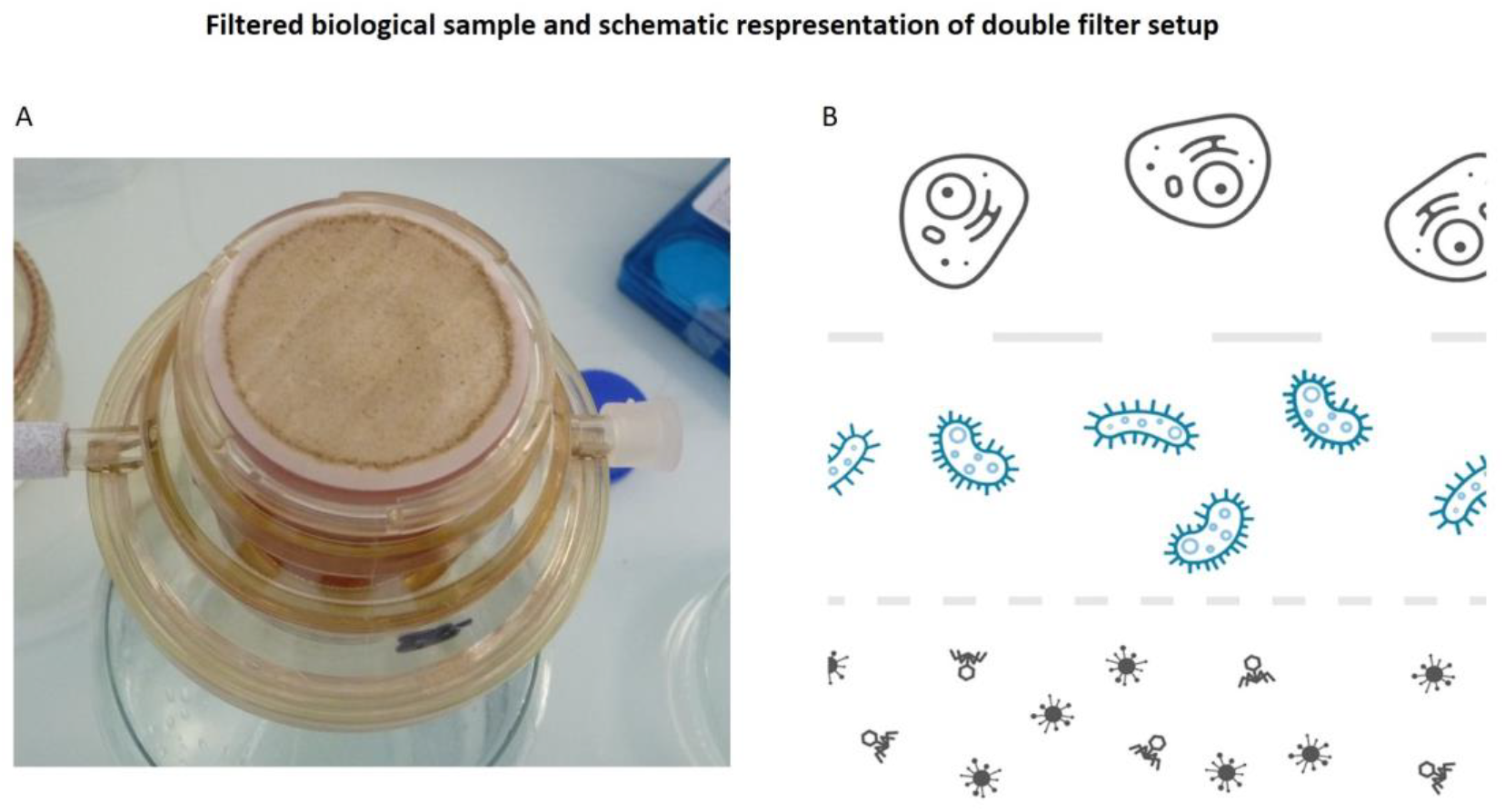
**A)** Filter setup; 0.22 μm containing biological material that represents the oceanic microbiome **B)** A schematic visualization of double filter setup. Discard eukaryotic cells during the first and viral/ phage content during the second filtering round.

To obtain high quality DNA we used the DNeasy PowerWater Kit (Qiagen), with minimal adjustments, according to the manufacturer’s protocol. The largest adjustment was supplementing an enzyme set (Lysozyme, Mutanolysin and Lysostaphin) for a more extensive cell lysis. DNA from both North Sea samples was sequenced subsequent to DNA isolation, however we obtained a suboptimal yield from DNA isolation of sample 2 and amplified the isolation to meet the minimal input requirements for sequencing. Sample 1 was filtered through the double filter setup and temporarily stored at −20 °C and long term stored at −80 °C, DNA isolation and sequencing were performed after approximately 11 months of storage.

### DNA library preparation, sequencing, data quality control and statistics

We used R9.4 flow cells for sequencing all three seawater samples. Libraries were prepared using rapid kits (SQK-RAD004) available at that time according to the manufacturer’s protocols (Oxford Nanopore Technologies, Oxford, UK). Data acquisition and base-calling were performed by MinKNOW (v19.06.8) controlling the MinION that sequenced the samples in 48-hours. Read-length and read-quality distributions were visualized using NanoPlot **[23]**, and read counts, base counts and average read lengths were obtained using custom made scripts.

### Using in silico PCR analysis to verify microbial genomes

To highlight the presence of microbial genomes FastPCR **[24]**was used to perform in silico PCR analysis using primer pair sequences for identification of bacteria and archaea. FastPCR allows users to upload a set of primer sequences and reports, among others, positions and length of hits found on the input data. We used the currently ‘best available’ rRNA primer pair, primer 1 and 2 are 17 and 21 bp long, respectively, with a total amplicon size of 464 bp (primer 1: 5’-CCTACGGGNGGCNGCAG-3’, primer 2: 5’-GACTACNNGGGTATCTAATCC-3’). FastPCR verifies both forward and reverse primer sequences and due to the erroneous nature of our long read technology we have set a threshold of =>80% alignment identity to the primer sequences, with the exception that no errors may occur at the last position on the 3’ end of the primer sequences. Since OneCodex is primarily tailored to classification of microbial data we used FastPCR, in a similar fashion, to verify any remaining microbial content in the unclassified reads.

### K-mer based metagenome characterization of microbial sequences from seawater

OneCodex uses a k-mer based taxonomic classification algorithm to characterize microbial data. It uses a reference database containing 53,193, 27,020, 1,724, 1,756 and 168 bacterial, viral, fungal, archaeal and protozoan genomes, respectively. A default k-mer size of 31 bp is used to break up every read from the input data and compares them to a database that contains every k-mer that is uniquely linked to a taxonomic group. OneCodex classifies reads based on a set of k-mers that together uniquely identify taxonomic groups, single read hits are taken as the minimum threshold for identification in this study. OneCodex also provides reads for which no unique taxonomic classification could be found, we subtracted these reads from the initial input data using the command line interface (CLI) provided by OneCodex. We filter these reads using a project ID (provided by the web interface), the original dataset and set the taxonomic label to 0. For these reads we inspect the presence of microbial 16S rDNA and repetitive content in an attempt to explain the unresolved classification.

### Assembly of long read metagenomics samples using the Flye assembler

Flye **[27]** is currently one of the few de novo assembly pipelines that allows genomic reconstruction of complex metagenomics samples with coverage as low as 2x. We have downloaded the assembly software from the GitHub repository (v2.6), used the metagenome default settings and provided the raw sequencing data. For sample 1 and 3 we used all available raw sequencing data, for computational effectiveness we used half of the sample 2 data set. We have verified the top-3 longest contigs using BLAST alignment with high homology parameters and selected the best hits based on largest query coverage with highest identity and literature references.

### Repetitive content analysis for unclassified reads

To investigate the repetitive nature of reads that remained unclassified after OneCodex characterization we used Tandem Repeat Finder software (v4.09) **[22]**, developed by Boston University, with default settings. The software locates repetitive patterns and reports their locations, sizes and copy numbers in a repeat table format. We have parsed both raw sequencing and unclassified data from sample 1, 2 and 3 to Tandem Repeat Finder and inspected the repeat occurrences on every read. With a custom-made script the frequencies of these occurrences on every read for every sample are summarized and expressed in 1, 2-5, 5-10 and >10 occurrences bins and plotted with R ggplot2 **[34]**.

## DISCUSSION

In this study, we have investigated the use of Nanopore sequencing for seawater metagenomics. Our main aims were to investigate the effectiveness of DNA isolation from samples directly obtained from the environment, optimize laboratory protocols for maximum sequencing results and evaluation of current metagenomics identification and assembly software. We used multiple isolation procedures, several different storage methods and subjected the data to a set of different analysis software. With only three ONT flow cells we were able to identify thousands of organisms, including bacteriophages, from which a large part at species level. It was possible to assemble genomes from environmental samples with Flye. In several cases this resulted in >1 Mbp contigs and in the particular case of a Thioglobus singularis species it even produced a near complete genome.

Although the enzyme cocktail used for cell lysis in our study was designed to break down cell walls for a wide range of bacteria there are potentially microbes that are immune to our lysis step. This might result in an underrepresentation of specific microbial communities compared to what truly thrives at these locations at that time. A possible solution, instead of lysing microbes with an enzyme set, would be to subject samples to mechanical lysis using silica beads or a combination of both. During experimental 12-hour sequencing runs (data not shown) we have observed that combining silica beads and enzymes during isolation yields significantly more sequencing data compared to isolation using only enzymes.

The double filter method separates eukaryotes and phages/viruses from bacteria in our sample. However, OneCodex still classifies a few hundred reads as either eukaryotic or viral. Eukaryota are particularly enriched for Dikarya, a subkingdom of fungi that are known to dominate the marine fungi fraction of environmental samples at European coastal regions **[32]**. These reads might have come from eukaryotic cells that are smaller than our largest filter (1.2 μm) or particles of these species that simply float around and were picked up by the smallest filter.

DNA molecules of our samples possibly suffered from fragmentation due to ice crystal formation during eleven months −20 and −80 °C storage. Additionally the yield of some sequencing runs is relatively low since biological material was dry frozen to the filter, making it more difficult to suspend the material during cell lysis. Under ideal circumstances DNA should be sequenced immediately after isolation circumventing DNA strand damage and loss of material.

The presence of viral DNA might be an indication that we have used too much water on a single filter, causing the accumulation of biological material to the point where the filter became saturated. A saturated filter might catch particles smaller than the smallest filter size and contaminate the isolation with material that would have otherwise passed through. On the other hand, viral DNA could be present due to infection of microbes, which could be recognized by inspecting flanking regions of the read containing the viral DNA to contain microbe specific genes. Additionally viruses could enter the microbial metagenomics pool when they are present at the outside of bacteria and pass through the double filter setup via hitchhiking.

We initially performed de novo assembly in order to find out whether we could obtain longer contigs for particularly abundant species. Due to low coverage and the high diversity in our sample it is no surprise that this was possible for just a limited number of species. It is actually encouraging that with such diversity and limited sequence depth we could still identify more than thousand organisms at the species level. Moreover, metagenomics analysis is a relative newcomer in the field of genetics hence both laboratory protocols and analytical pipelines still need improvement to result higher accurate and more robust solutions for sample such as seawater.

We have performed an in silico PCR analysis to identify 16S RNA sequences in our raw sequence dataset. This showed that even under high error rate conditions reads contain enough homology to detect well conserved genes. This method could potentially be utilized to detect other genes in a similar fashion, for example genes that encode antibiotics biosynthesis.

While OneCodex was able to identify the diversity of a substantial amount of our samples, it could not resolve any classification for a large part of our data. The large k-mer size is most probably a crucial factor for unclassified data, due to the relatively low quality (approximately 10% error) of long-read data 10 bp would be a more suitable k-mer size. We confirmed that the data quality of these reads (both read length and quality distributions) are within acceptable bounds and observed no particular repetitive element enrichment compared to the reads that contributed to classifications.

Sequences that are representative of species that are currently unknown might explain the unclassified state of those reads, and are therefore valuable for contribution of a deeper understanding of the microbial marine fingerprint. Moreover, open access databases might not contain genomic information on particular microbes since obtaining genomes that are particularly large or come from non-culturable (non-culturable organisms are indicated with ‘Candidatus’ labels) microbes remains a complicated task. For example, although available, protists are poorly represented in the OneCodex database, perhaps because their genomes are often extremely large (for dinoflagellates up to 270 Gbp **[33]**). Hence, microbes that are less thoroughly investigated might not have been included into the OneCodex genome selection. Since OneCodex is tailored to the identification of single cell organisms it probably will leave reads from multicellular organisms unclassified. Although no strong evidence was observed, lenient BLAST alignment of the top-3 longest reads of every sample did identify some small homologue regions with sequences from plant or algae in the NCBI database.

Despite the fact that these experiments are pilot studies we have observed promising results for both laboratory protocols and species identifications analysis. As described above, sample collection, DNA isolation and species identification is still hindered by both technical and biological difficulties. However our method provides a good impression on the elegance of our method that comes from its robustness and simplicity. We have performed equivalent experiments in student field practical assignments with similar marine samples, and students showed that even under more restricted conditions (12-hour sequencing runs) large biodiversity could still be detected. This indicates that the simplicity of our setup provides an ideal setting for student exercises, that will surely facilitate educational programs in genetics and bioinformatics.

## DATA AVAILABILITY

Data submission in process and will be available at NCBI. The data is temporarily available at: https://mliem.stackstorage.eom/s/i1XQRymbs1xSQk4

## DECLARATIONS

Ethics approval and consent to participate - not applicable

Consent for publication - not applicable

Competing interests - authors declare no competing interest

Funding - No external funding

Authors’ contributions - all authors contributed to the writing of the manuscript, ML performed bioinformatics analysis, AJGRT and ML performed lab experiments, ML and HPS wrote the first draft of the manuscript. CVH and HPS supervised the study.

## ACKNOWLEDGEMENT

We would like to express our gratitude to OneCodex for answering questions on the available genome selection and the help with the CLI, and Future Genomics Technologies (Leiden) for the help with initial sequencing runs. All authors contributed equally to the manuscript.

## REFERENCES

1. Zobell, C. E., Marine Microbiology, Chronica Botanica Co, Waltham, Mass., USA, 1946, p. 240.

2. Velankar, N. K., Bacteria isolated from seawater and marine mud off Mandapam (Gulf of Mannar and Palk Bay). Indian J. Fish., 1957, 4, 208–227.

3. Wood, E. J. F., Some aspects of marine microbiology. J. Mar. Biol. Assoc. India, 1959, 1, 26–32.

4. Marine microbial diversity. https://www.ncbi.nlm.nih.gov/pubmed/28586685

5. The Sorcerer II Global Ocean Sampling expedition: northwest Atlantic through eastern tropical Pacific. https://www.ncbi.nlm.nih.gov/pubmed/17355176/

6. Marine Rare Actinobacteria: Isolation, Characterization, and Strategies for Harnessing Bioactive Compounds (review) https://www.ncbi.nlm.nih.gov/pmc/articles/PMC5471306/

7. Metagenomics uncovers a new group of low GC and ultra-small marine Actinobacteria. https://www.ncbi.nlm.nih.gov/pubmed/23959135

8. The Tara Pacific expedition-A pan-ecosystemic approach of the “-omics” complexity of coral reef holobionts across the Pacific Ocean https://www.ncbi.nlm.nih.gov/pubmed/31545807

9. Characterization of the microbial community diversity and composition of the coast of Lebanon: Potential for petroleum oil biodegradation https://www.ncbi.nlm.nih.gov/pubmed/31425842

10. Depth and location influence prokaryotic and eukaryotic microbial community structure in New Zealand fjords https://www.ncbi.nlm.nih.gov/pubmed/31377366

11. High-throughput sequencing and analysis of microbial communities in the mangrove swamps along the coast of Beibu Gulf in Guangxi, China. https://www.ncbi.nlm.nih.gov/pubmed/31253826

12. Environmental Genome Shotgun Sequencing of the Sargasso Sea https://www.ncbi.nlm.nih.gov/pubmed/15001713

13. Metagenomics of the deep Mediterranean, a warm bathypelagic habitat. https://www.ncbi.nlm.nih.gov/pubmed/17878949/

14. High resolution profiling of coral-associated bacterial communities using full-length 16S rRNA sequence data from PacBio SMRT sequencing system https://www.ncbi.nlm.nih.gov/pubmed/28584301

15. Influence of 16S rRNA variable region on perceived diversity of marine microbial communities of the Northern North Atlantic. https://www.ncbi.nlm.nih.gov/pubmed/31344223

16. Metagenomics uncovers a new group of low GC and ultra-small marine Actinobacteria https://www.ncbi.nlm.nih.gov/pmc/articles/PMC3747508/

17. Genome Streamlining in a Cosmopolitan Oceanic Bacterium https://science.sciencemag.org/content/309/5738/1242.long

18. Time-series analyses of Monterey Bay coastal microbial picoplankton using a ‘genome proxy’ microarray. https://www.ncbi.nlm.nih.gov/pubmed/20695878

19. Proteorhodopsin photosystem gene clusters exhibit co-evolutionary trends and shared ancestry among diverse marine microbial phyla https://www.ncbi.nlm.nih.gov/pubmed/17359257

20. Proteorhodopsin genes are distributed among divergent marine bacterial taxa. https://www.ncbi.nlm.nih.gov/pubmed/14566056

21. Genomic content of uncultured Bacteroidetes from contrasting oceanic provinces in the North Atlantic Ocean. https://www.ncbi.nlm.nih.gov/pubmed/21895912

22. G. Benson,“Tandem repeats finder: a program to analyze DNA sequences” Nucleic Acids Research (1999) Vol. 27, No. 2, pp. 573–580.

23. https://github.com/wdecoster/NanoPlot

24. Kalendar R, Khassenov B, Ramankulov Y, Samuilova O, Ivanov KI 2017. FastPCR: an in silico tool for fast primer and probe design and advanced sequence analysis. Genomics, 109: 312–319. DOI: 10.1016/j.ygeno.2017.05.005

25. Evaluation of general 16S ribosomal RNA gene PCR primers for classical and next-generation sequencing-based diversity studies https://www.ncbi.nlm.nih.gov/pmc/articles/PMC3592464/

26. One Codex: A Sensitive and Accurate Data Platform for Genomic Microbial Identification Samuel S Minot, Niklas Krumm, Nicholas B. GreenfieldPublished 2015 DOI:10.1101/027607

27. Kolmogorov M. et al. (2018) Assembly of Long Error-Prone Reads Using Repeat Graphs. https://doi.org/10.1093/bioinformatics/bty956.

28. Genome Sequence of “Candidatus Thioglobus singularis” Strain PS1, a Mixotroph from the SUP05 Clade of Marine Gammaproteobacteria. https://www.ncbi.nlm.nih.gov/pubmed/26494659

29. Abundant SAR11 viruses in the ocean. https://www.ncbi.nlm.nih.gov/pubmed/23407494

30. Marine prasinovirus genomes show low evolutionary divergence and acquisition of protein metabolism genes by horizontal gene transfer. https://www.ncbi.nlm.nih.gov/pubmed/20861243

31. Complete Genome Sequence of the Fish Pathogen Flavobacterium columnare Strain C#2 https://www.ncbi.nlm.nih.gov/pubmed/27340080

32. Molecular diversity and distribution of marine fungi across 130 European environmental samples https://royalsocietypublishing.org/doi/10.1098/rspb.2015.2243

33. The exceptionally large genome of the harmful red tide dinoflagellate Cochlodinium polykrikoides Margalef (Dinophyceae): determination by flow cytometry, Algae 2016; 31(4): 373–378, DOI: https://doi.org/10.4490/algae.2016.31.12.6

34. H. Wickham. ggplot2: Elegant Graphics for Data Analysis. Springer-Verlag New York, 2016

